# Metabolic reprogramming and gut microbiota ecology drive divergent *Plasmodium vivax* infection outcomes in *Anopheles darlingi*

**DOI:** 10.1101/2025.08.13.670040

**Authors:** Bianca Cechetto Carlos, Kamila Voges, Pedro Henrique de Andrade Affonso, Amie Jaye, Carlos Tong Rios, Bruno Tinoco-Nunes, Diego Peres Alonso, Robert M. MacCallum, Marta Moreno, Dina Vlachou, Jayme A. Souza-Neto, George K. Christophides

## Abstract

*Anopheles darlingi* is the principal malaria vector in the Amazon basin, where *Plasmodium vivax* accounts for the majority of cases. Despite its epidemiological importance, the molecular and microbial determinants of *A. darlingi* susceptibility to *P. vivax* remain poorly understood. Here, we investigated vector-parasite-microbiota interactions using experimental infections with field-derived *P. vivax* gametocytaemic blood, which produced two distinct infection phenotypes: low and high oocyst burdens. Transcriptomic profiling of mosquito midguts across key parasite developmental timepoints revealed that low-infection mosquitoes mounted an early and sustained response characterised by activation of detoxification pathways, redox regulation, aromatic amino acid catabolism, and purine depletion, likely coordinated through neurophysiological cues, which collectively create a metabolically restrictive environment for parasite development. These physiological changes were accompanied by reduced bacterial diversity and enrichment of Enterobacteriales and Pseudomonadales, taxa previously linked to anti-*Plasmodium* activity. Conversely, high-infection mosquitoes exhibited limited metabolic reprogramming, expansion of Flavobacteriales, and transcriptional signatures consistent with permissive physiological states, potentially associated with reproductive trade-offs. Importantly, low infection outcomes consistently arose from bloodmeals with the lowest gametocyte densities, suggesting that host- and parasite-derived components of the bloodmeal act as early conditioning factors that prime the mosquito midgut for either resistance or susceptibility. These findings reframe *A. darlingi* vector competence to *P. vivax* not as a fixed immune trait but as a dynamic outcome of early redox, metabolic, and microbial interactions. They also highlight ecological and physiological targets for transmission-blocking strategies and reinforce the importance of studying vector-parasite interactions in regionally relevant systems.

## Introduction

Malaria, a mosquito-borne disease caused by protozoan parasites of the genus *Plasmodium*, remains a major global health challenge. In 2023, an estimated 263 million people were infected, resulting in 597,000 deaths worldwide [1]. While *Plasmodium falciparum* accounts for most global malaria cases and fatalities, *Plasmodium vivax* continues to cause a substantial proportion of the burden, particularly outside sub-Saharan Africa. In the Americas, *P. vivax* is the predominant species, responsible for 68-72% of reported malaria cases, with the highest burden concentrated in countries of the Amazon basin. Transmission in this region is largely confined to rural and remote communities, where limited access to healthcare, diagnostics, and treatment impedes control efforts [2]. The ability of *P. vivax* to relapse via dormant liver-stage hypnozoites presents an additional challenge, sustaining transmission even in areas of low endemicity and complicating elimination strategies.

*Anopheles darlingi* is the principal malaria vector in the Amazon basin. This neotropical mosquito species is highly anthropophilic, endophilic, aggressive, opportunistic and susceptible to *Plasmodium* infection [3–5]. Despite its critical epidemiological importance in the Americas, *A. darlingi* remains largely understudied, and the molecular mechanisms underpinning its interactions with *Plasmodium* are still poorly understood.

After ingestion by female mosquitoes during a blood meal, *Plasmodium* gametocytes rapidly differentiate into gametes in the mosquito midgut. Fertilization leads to zygote formation, which subsequently develops into motile ookinetes. These ookinetes traverse the peritrophic matrix and the midgut epithelium, eventually anchoring beneath the basal lamina to form oocysts. The traversal of the midgut, typically between 18 and 26 hours post-bloodmeal, represents a major population bottleneck for the parasite [6–8]. Parasites that survive this barrier multiply within oocysts via sporogony, producing thousands of sporozoites. Between 9 and 12 days later, mature oocysts rupture, releasing sporozoites into the hemocoel, from where they migrate to the salivary glands for transmission to a new vertebrate host.

The successful completion of this complex *Plasmodium* developmental journey is shaped by multiple vector-derived factors, particularly the mosquito innate immune response and midgut microbiota composition [9–11]. Genome-wide transcriptomic studies and functional analyses in model systems have uncovered a diverse immune landscape that governs mosquito susceptibility to infection [12–14]. These immune responses are not only responsive to *Plasmodium* but also to bacteria, indicating extensive crosstalk between antibacterial and antiparasitic defenses [15–18].

Our current understanding of these immune responses is almost entirely based on studies of *A. gambiae* infected with *P. falciparum* or rodent parasites such as *P. berghei*. In contrast, the *A. darlingi-P. vivax* transmission system, which is central to malaria persistence in the Americas, has been largely neglected. This knowledge gap stems in part from the relatively late development of genomic and experimental resources: the first draft genome of *A. darlingi* was published only in 2013 [19], and stable laboratory colonies were established in Peru, Brazil and French Guiana within the last decade [20–23]. A more fundamental barrier is the absence of a continuous in vitro culture system for *P. vivax* [24], which restricts experimental infection studies to field settings with simultaneous access to both laboratory-reared mosquitoes and blood from naturally infected gametocyte carriers [3, 22, 25].

Here we investigate the tripartite interaction between *A. darlingi*, its midgut microbiota, and *P. vivax* using a semi-field experimental system in Iquitos, Peru. By infecting colony-reared *A. darlingi* with blood from naturally infected *P. vivax* carriers and combining transcriptomic and microbiome analyses across multiple timepoints and infection intensities, we identify metabolic, and microbiota trajectories associated with vector competence. Our findings suggest that *P. vivax* developmental success is shaped by orchestrated transcriptional reprogramming of mosquito metabolism and shifts in microbiota diversity, likely triggered by host- and/or parasite-derived factors in the bloodmeal. These findings provide new insight into a neglected but epidemiologically important malaria transmission system.

## Results & Discussion

### Experimental infections reveal distinct oocyst load phenotypes

We conducted experimental *A. darlingi* infections using blood samples from six *P. vivax* gametocytaemic patients (P1-P6) with varying gametocyte densities (**Dataset 1A**). Adult female *A. darlingi* mosquitoes were divided into 12 groups: six were fed with untreated blood containing potentially infectious gametocytes (I groups), and six with heat-inactivated blood serving as non-infectious controls (C groups) using direct membrane feeding assays (DMFAs).

Oocyst enumeration in dissected midguts 7 days post blood feeding (pbf) revealed a complete absence of infection in all C groups, validating the heat inactivation protocol [26]. In contrast, all I groups displayed varying degrees of infection (**Fig. 1A** and **Dataset 1B**). Two distinct infection phenotypes emerged: a low oocyst burden group (P1, P3, P4) and a high oocyst burden group (P2, P5, P6). Median and mean oocyst loads were largely concordant across groups, except for P6, suggesting a near-normal distribution of infection intensities in most groups. This contrasts with the commonly observed overdispersion in *P. falciparum* infections of *A. gambiae* [13, 26].

**Figure 1.**
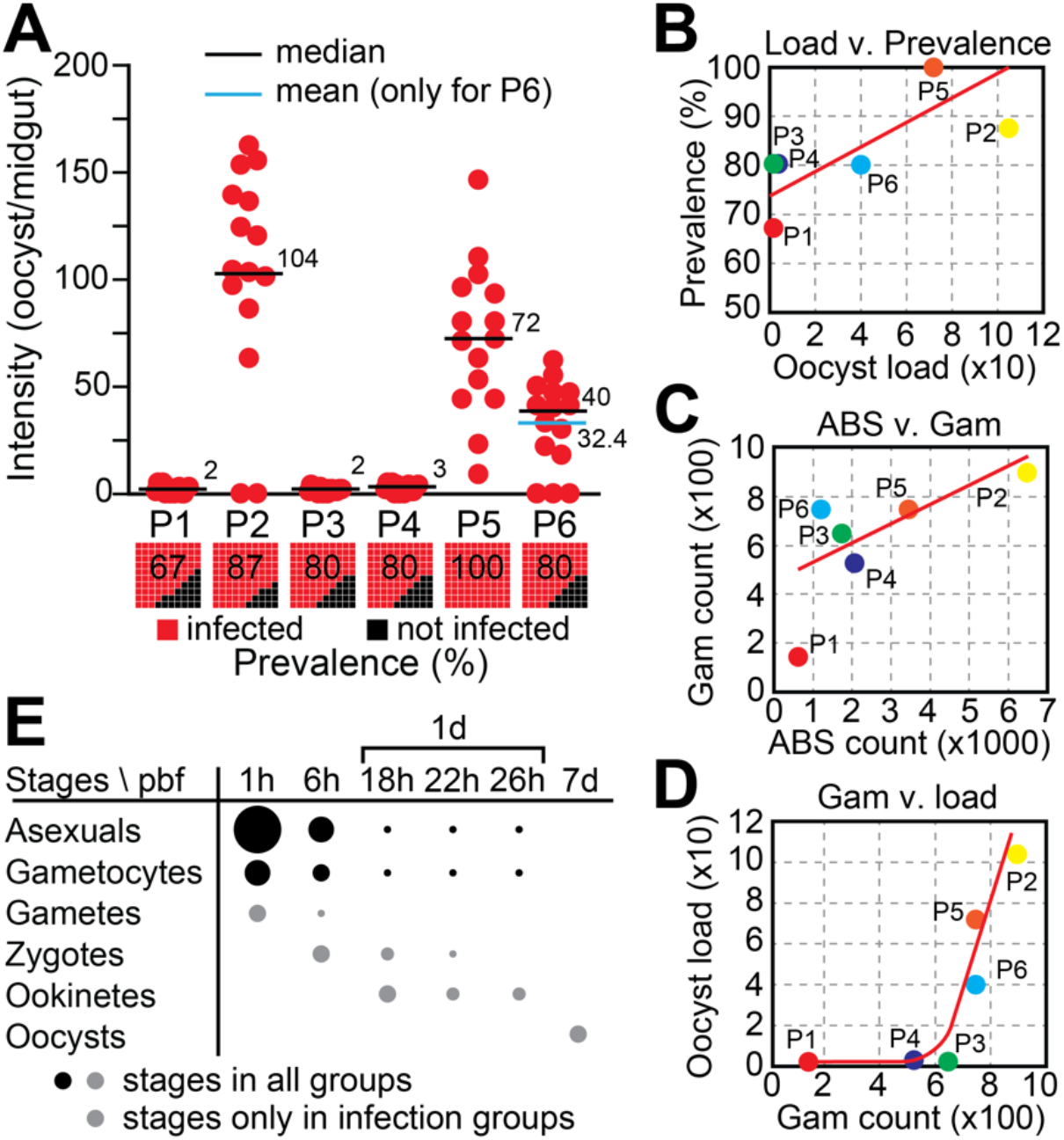
Experimental infection of *A. darlingi* with *P. vivax* field isolates. **(A)** Oocyst load per midgut in mosquitoes fed on blood from six gametocytaemic patients (P1-P6). Each red dot represents a single midgut from an infected mosquito; black squares represent uninfected individuals. Horizontal lines indicate median oocyst load (black); the mean is also shown for P6 (blue). Numbers above bars indicate median values. Bar plots below show infection prevalence (% infected, red; % uninfected, black) in each group. **(B)** Positive correlation between oocyst load and infection prevalence across experimental groups; red line indicates linear regression. **(C)** Correlation between ABS counts and gametocyte densities in patient blood samples. **(D)** Positive correlation between gametocyte density and oocyst load. A gametocyte threshold of ~700/µL appears necessary for high infection intensities. **(E)** Expected presence of parasite developmental stages in the mosquito midgut over time post blood feeding (pbf). Sampling timepoints: 1h, 6h, 18h, 22h, 26h, and 7d pbf. Dot size represents relative abundance; black dots indicate stages expected across all groups, grey dots those expected only in infected mosquitoes.

Interestingly, oocyst prevalence remained high and relatively uniform across infection groups (67%-100%), regardless of the oocyst burden (**Fig. 1B**). This decoupling of infection prevalence from intensity suggests that factors beyond simple gametocyte presence influence oocyst load. Notably, gametocyte density positively correlated with the abundance of asexual blood stages (ABS) (**Fig. 1C**), and high infection loads were only observed when gametocyte counts exceeded ~700/µL of blood (**Fig. 1D**), suggesting a possible threshold effect in infectivity.

To capture key developmental stages of *P. vivax* within the mosquito, midguts were sampled at six timepoints post blood feeding: 1h (hour), 6h, 18h, 22h, 26h, and 7d (days) pbf (**Fig. 1E**). Based on established timelines of parasite development, we expect that at 1h pbf the midguts would contain ABS, gametocytes, and newly emerged gametes. By 6h pbf, zygote presence is anticipated, while ookinetes are expected to appear between 18h and 26h pbf. By 7d pbf, only mature oocysts are expected to remain, following the digestion of the blood bolus and progression of parasite development.

### Transcriptional profiling highlights infection-specific gene expression programmes

A total of 72 RNA-seq libraries were generated from *A. darlingi* midguts across the six timepoints pbf, spanning high and low *P. vivax* infection groups and their respective controls. Sequencing yielded an average of 15.5 million reads per library for low infections and 34.5 million reads for high infections. About 60% of reads mapped to the *A. darlingi* reference genome AdarC3, identifying transcripts corresponding to 10,895 annotated genes: 10,423 with *A. gambiae* orthologs and 472 with no identifiable *A. gambiae* orthologs.

Unsupervised hierarchical clustering based on global transcriptome similarity revealed consistent temporal organization among samples (**Fig. 2A**). For high infection groups, samples from 18h, 22h, and 26h pbf clustered tightly. These timepoints were therefore grouped together and referred to as “1d pbf” in subsequent analyses. Notably, within this 1d pbf group, I and C samples separated into distinct clusters, indicating that *P. vivax* infection elicits specific transcriptional responses. A similar pattern was observed in low infection groups, although the separation between I and C samples at 1d pbf was less pronounced. This suggests that the mosquito transcriptional response is infection-dependent and potentially modulated by the magnitude or quality of the infecting parasite population.

**Figure 2.**
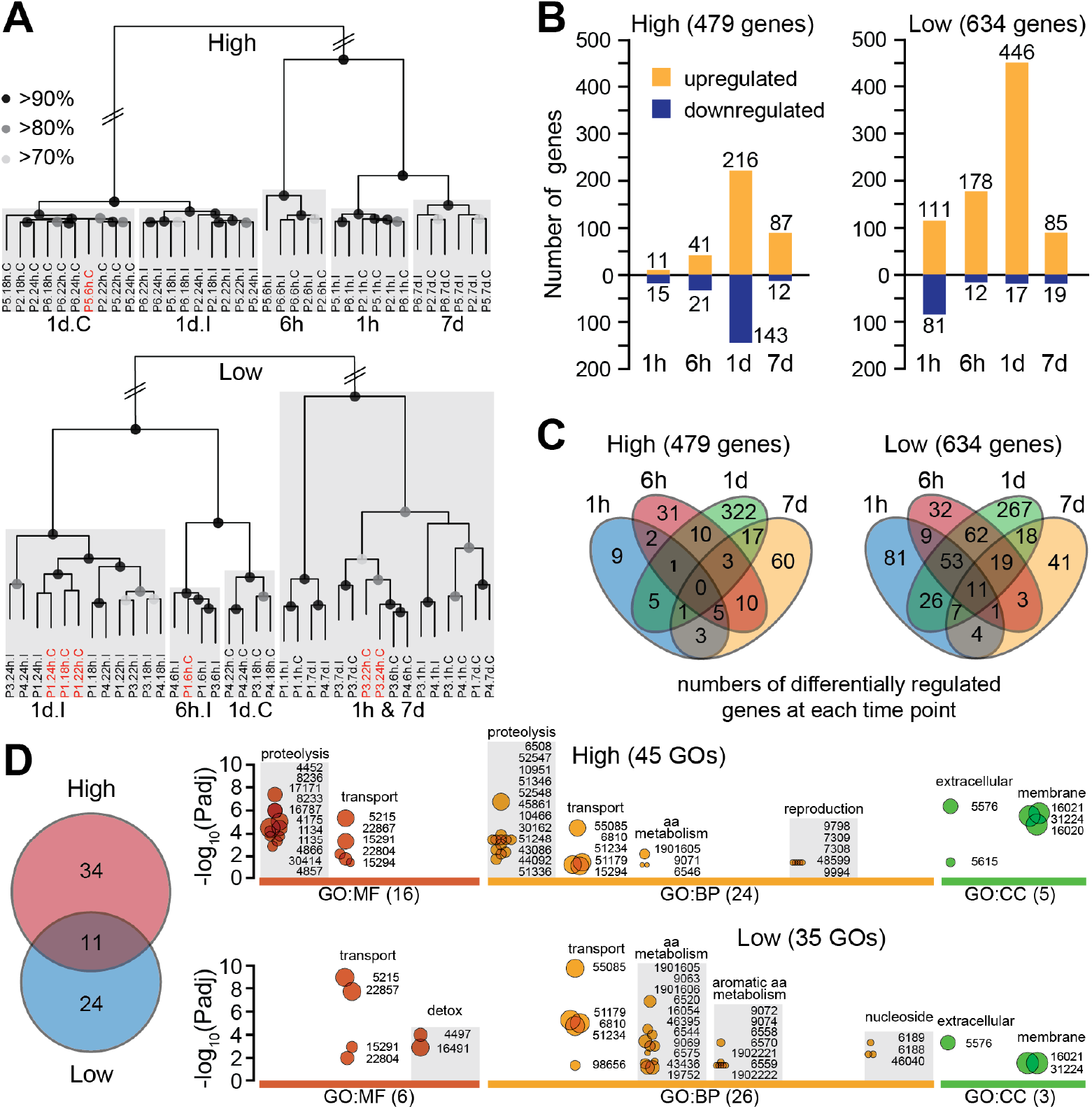
Differential transcriptional responses in *A. darlingi* midguts during high and low *P. vivax* infections. **(A)** Hierarchical clustering of RNA-seq libraries from *A. darlingi* midguts sampled at multiple timepoints (1h, 6h, 18h, 22h, 26h, 7d) post blood feeding. Samples from high (top panel) and low (bottom panel) infection groups are shown separately. Grey boxes indicate clusters by timepoint; black dots on branches denote bootstrap support levels: >90% (large), >80% (medium), >70% (small). **(B)** Number of genes differentially regulated (infected vs. control) across timepoints in high (left) and low (right) infection groups. Orange bars: upregulated genes; blue bars: downregulated genes. **(C)** Venn diagrams showing overlap of differentially regulated genes across four timepoints (1h, 6h, 1d, 7d) in high (left) and low (right) infection groups. **(D)** Gene ontology (GO) enrichment analysis of DEGs from the same four timepoints. The Venn diagram indicates 11 GO terms shared between infection groups, with 34 and 24 terms uniquely enriched in high and low infection groups, respectively. Bubble plots show significantly enriched GO terms (Padj < 0.05) in three GO categories: molecular function (MF), biological process (BP), and cellular component (CC). Bubble size reflects the number of associated genes, ranging from 3 (smallest) to 177 (largest).

Differential gene expression analysis revealed 479 (**Dataset 2**) and 634 (**Dataset 3**) differentially expressed genes (DEGs) in high and low infection groups, respectively (**Fig. 2B**). The majority of DEGs were detected at 1d pbf in both cases (359 in high and 463 in low infections), coinciding with the critical window of midgut traversal. Whereas DEGs at this timepoint were moderately balanced between upregulated and downregulated genes in high infection groups, 96.3% of DEGs in low infection groups were upregulated, indicating a strong transcriptional response that may play a role in limiting infection. Earlier in the infection course, high infection groups exhibited a muted response, with only 11 and 41 upregulated DEGs at 1h and 6h pbf, respectively, compared to 111 and 178 in the corresponding low infection groups.

This early and sustained activation in low infection groups suggests a more rapid and effective response that may hinder parasite infection. In support of this, Venn diagram analysis of DEGs across timepoints revealed a substantial overlap in the low infection group between genes upregulated at 1h, 6h, and 1d pbf (**Fig. 2C**), consistent with a primed and coordinated early response.

### Metabolic deprivation and redox regulation define divergent infection trajectories

A comparative analysis of gene ontology (GO) enrichment in highly versus lowly infected mosquito groups revealed clear differences in the host response, reflecting divergent physiological engagements. A total of 45 significantly enriched GO terms were identified in the high and 35 in the low infection groups, spanning molecular function (MF), biological process (BP), and cellular component (CC) categories (**Fig. 2D** and **Dataset 4**). Notably, 11 GO terms were shared between the two groups, encompassing terms such as transporter activity and alpha-amino acid metabolism. This overlap likely represents core physiological responses to infection, involving solute exchange and protein turnover, irrespective of infection intensity.

The high infection group was uniquely enriched for GO terms related to proteolysis and its regulation, as well as pathways linked to oocyte development. As there is significant overlap between genes involved in immune responses and proteolysis, such as clip-domain serine proteases and their homologs, this pattern could suggest a balanced response to the parasite challenge coupled with a trade-off in reproductive investment (**Fig. 2D**). In contrast, mosquitoes in the low infection group exhibited a distinct response dominated by genes involved in alpha and aromatic amino acid catabolism, including phenylalanine and tyrosine, as well as purine biosynthesis. These metabolic processes are particularly relevant given that both aromatic amino acids and purines are essential for *Plasmodium*, which lacks the biosynthetic capacity to produce them [27, 28]. This suggests that the mosquito may limit parasite development through metabolic deprivation or sequestration of these critical nutrients.

Additionally, the enrichment of monooxygenase and oxidoreductase activity in the low infection group suggests a shift toward redox regulation and detoxification pathways (**Fig. 2D**), possibly linked to elevated mitochondrial activity or microbial dysbiosis. Together, these findings support the view that midgut transcriptional responses related to the mosquito metabolic state may actively determine the trajectory of *P. vivax* infection.

### Time-course analysis identifies early divergence in infection trajectories

To further dissect the mechanisms underlying the divergent *P. vivax* infection outcomes, we extended our analysis beyond GO enrichment by examining the temporal dynamics of DEGs at each of the four timepoints pbf (**Dataset 5**). This high-resolution analysis allowed us to pinpoint specific transcriptional changes that may influence the final infection outcome.

Interestingly, temporal correlation analysis between high and low infection groups revealed striking divergence at the earliest (1h) and most critical (1d) timepoints (**Fig. 3**). At 1h pbf, responses were largely uncorrelated, reflecting highly distinct early physiological states between mosquitoes that would go on to develop either high or low parasite burdens (Pearson r = 0.03, R^2^ = 0.0009, p = 0.67). A similar lack of correlation was observed at 1d pbf, coinciding with the peak of microbial load and activity in the midgut and the critical window of establishment of infection, suggesting that transcriptional trajectories diverge most sharply at times when parasite establishment is most vulnerable to vector responses (Pearson r = –0.07, R^2^ = 0.0049, p = 0.063). In contrast, at 6h pbf, gene expression patterns between high and low infection groups showed greater convergence, suggesting transient homeostatic realignment after the initial response to blood feeding. By 7d pbf, when the parasites have matured into oocysts and midgut activity has stabilised, expression profiles again became more correlated, consistent with a return to physiological equilibrium.

These findings highlight the 1h and 1d pbf in the low infection group as the most informative timepoints for understanding the molecular drivers of infection outcome. The early divergence at 1h likely reflects immediate transcriptional reprogramming triggered by bloodmeal composition and, possibly, microbiota response, while the divergence at 1d pbf coincides with the outcome-determining phase of parasite midgut traversal and immune engagement. We therefore focused subsequent analyses on these two key timepoints to uncover mechanisms that underpin the establishment or restriction of infection.

**Figure 3.**
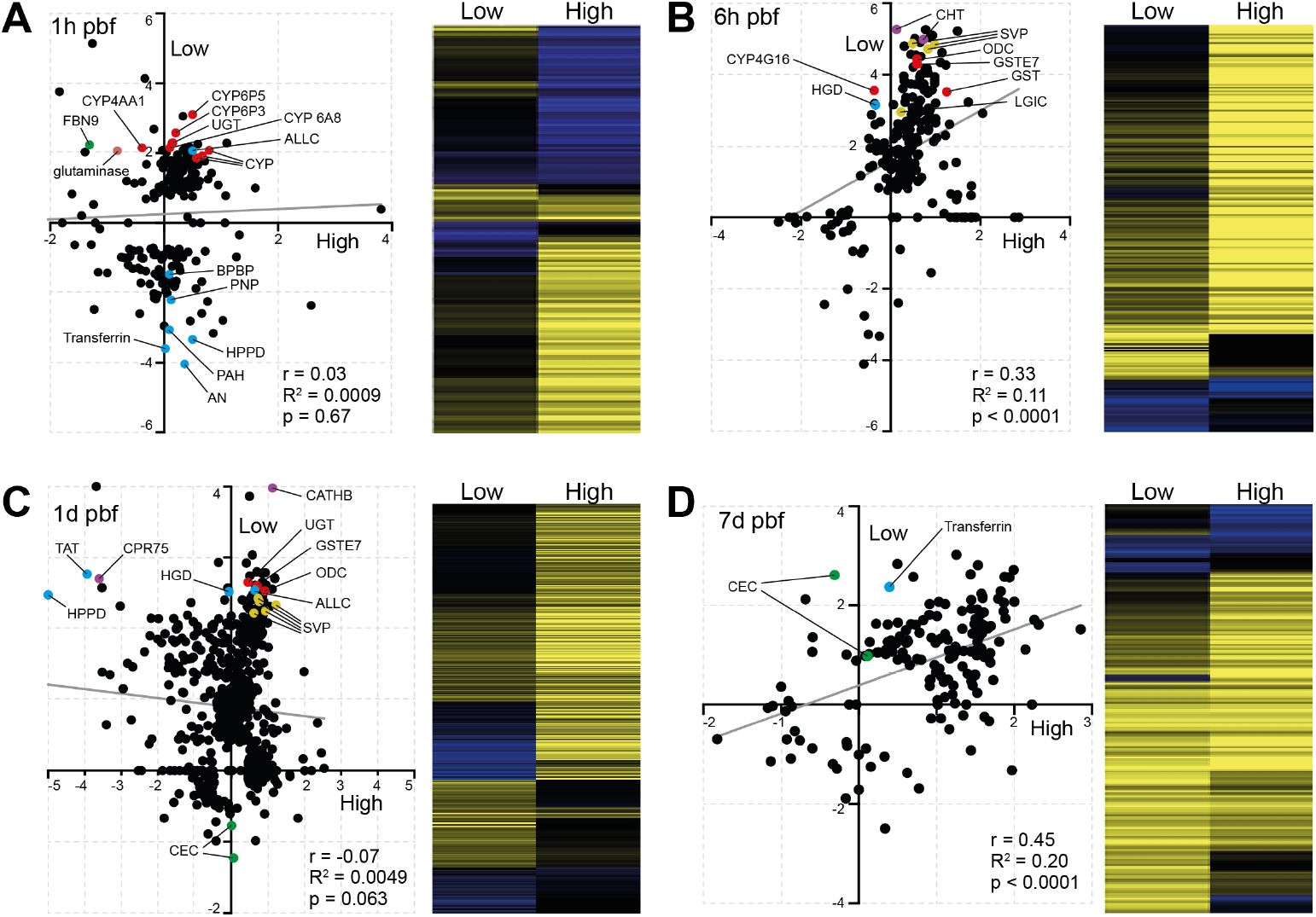
Temporal divergence of *A. darlingi* transcriptional responses resulting in high and low *P. vivax* infection outcomes. Scatter plots (left panels) compare log2 fold changes in gene expression (infected vs. control) between high and low infection groups at the four timepoints pbf: 1h **(A)**, 6h **(B)**, 1d **(C)**, and 7d **(D)**. Each point represents a DEG. Pearson correlation coefficient, R^2^ values, and p-values are shown within each panel. The least squares regression line in each scatter plot is shown in grey. Select DEGs discussed in the main text are highlighted in colour to reflect functional groupings: red, detoxification and redox regulation; blue, purine and amino acid metabolism; green, immune-related genes; yellow, stress or cell turnover-related genes; violet, others including neuronal and structural components. Heatmaps (right panels) display the corresponding expression profiles across conditions, with yellow indicating upregulation and blue indicating downregulation.

### Low infection is marked by early redox activation and metabolic reprogramming

At 1h pbf, mosquitoes in the low infection group displayed a striking upregulation of genes associated with detoxification and redox regulation (**Fig. 3A**). Multiple cytochrome P450s (e.g., CYP6P5, CYP6P3, CYP4AA1, CYP9b2) and glutathione S-transferases (e.g., GSTE2 and GSTE7) were upregulated, along with glutaminase, an enzyme that liberates glutamate and urea from glutamine. These changes indicate early oxidative stress responses and a detoxification state in the midgut.

Metabolic interference also emerged as a key theme. We observed upregulation of allantoicase (ALLC), a purine-degrading enzyme, along with concurrent downregulation of multiple genes involved in purine salvage and biosynthesis, such as purine nucleoside phosphorylase (PNP), phenylalanine-4-hydroxylase (PAH), 4-hydroxyphenylpyruvate dioxygenase (HPPD), and a bifunctional purine biosynthesis protein (BPBP). These shifts suggest programmed depletion of purine and nucleoside resources, which are essential for parasite DNA and RNA synthesis, especially given parasite reliance on host purines [28].

Further metabolic reprogramming was evident through the modulation of AMP-dependent CoA ligases, glucosyl/glucuronosyl transferases (UGT), and various solute transporters. Transferrin was strongly downregulated, hinting at reduced iron availability, while carbonic anhydrase suppression suggests altered pH regulation and gas exchange. Collectively, these responses point to a nutritionally restrictive and immunologically hostile midgut environment.

Interestingly, classical immune pathway components were not strongly activated at this stage, except for the fibrinogen-like recognition protein *FBN9*, which was significantly upregulated in low infection mosquitoes but downregulated in the high infection group. Given the known roles of FBN9 in both microbial recognition and immune modulation [29], this suggests early microbiota-driven immune priming may be linked to the low infection phenotype.

In contrast, the high infection group exhibited a muted transcriptional response at 1 h pbf, with only a small number of upregulated genes and minimal immune or metabolic reprogramming. This lack of early transcriptional activation may contribute to the permissive environment that allows *P. vivax* to proceed through midgut traversal and establish oocysts.

Together, these early timepoint comparisons provide mechanistic insight into how differential regulation of redox homeostasis, purine metabolism, and microbial recognition may drive the divergent *P. vivax* infection outcomes observed in *A. darlingi*. The early and sustained activation of these processes in low infection groups supports a model in which rapid physiological adaptation disrupts parasite development before midgut invasion is complete.

At 6h pbf, a moderate positive correlation in gene expression was observed between high and low infection groups (Pearson r = 0.33, R^2^ = 0.11, p < 0.0001), indicating partial convergence of transcriptional responses (**Fig. 3B**). This alignment may reflect a transient state of physiological homeostasis following the initial perturbation induced by blood ingestion. However, consistent with the patterns observed at 1h pbf, genes associated with redox regulation, including two GSTs and a cytochrome P450, remained strongly upregulated in the low infection group. In addition, homogentisate 1,2-dioxygenase (HGD), an enzyme involved in the degradation of aromatic amino acids tyrosine and phenylalanine, was also strongly upregulated. Given the importance of these amino acids for parasite survival and their role in the synthesis of folate and para-aminobenzoic acid (PABA) that are essential in nucleotide biosynthesis [30], HGD upregulation may reflect a host-driven strategy to deprive the parasite of essential metabolites. Additional genes involved in purine metabolism, such as amidophosphoribosyltransferase, were also upregulated though more moderately, further supporting the hypothesis that nutritional deprivation, particularly of purines and nucleotides, may underpin the low infection phenotype.

Among the most strongly upregulated genes (32-38-fold change) were two previously unannotated genes (ADAC003727, ADAC003402) involved in chitin metabolism. These likely relate to peritrophic matrix dynamics, possibly in response to microbial dysbiosis [31]. The strong upregulation of ornithine decarboxylase (ODC), a key enzyme in the biosynthesis of polyamines that are essential for the DNA/RNA stability, is also worth noting, as this may suggest disruption of gut epithelial homeostasis due to microbiota dysbiosis [32].

Intriguingly, several neuronal signalling components, including three synaptic vesicle protein (SVP) genes and a ligand-gated ion channel (LGIC), were also among the top DEGs in the low infection group. SVPs are involved in neurotransmitter storage and release, while LGICs mediate neuronal signal transduction. Their coordinated induction raises the possibility of a gut-neural axis response modulating midgut physiology or homeostasis in reaction to the infectious bloodmeal [33], potentially contributing to a more restrictive environment for parasite development.

### Sustained transcriptional activity and epithelial modulation during midgut invasion

The 1d (18-26h) pbf timepoint marked the peak of transcriptional activity in both high and low infection groups, as well as the point of greatest divergence between them (**Fig. 3C**). No correlation was observed in gene expression profiles (Pearson r = −0.07, R^2^ = 0.0049, p = 0.063), indicating that distinct transcriptional programs were engaged as the parasite attempted midgut invasion. This window corresponds with the peak of microbial proliferation and the critical stage of ookinete traversal through the midgut epithelium, when vector responses can decisively influence infection success [34, 35]. While no single dominant transcriptional pathway stood out, the patterns observed at earlier timepoints were broadly reinforced.

In the low infection group, genes associated with detoxification and cellular stress, including UGT, GSTE7 and ODC, remained strongly upregulated. This sustained stress response likely maintains an unfavourable physiological state for parasite development. Consistent with earlier stages, multiple enzymes linked to purine and aromatic amino acid degradation were also strongly upregulated.

These included ALLC, HGD, tyrosine aminotransferase (TAT), and HPPD, enzymes that sequentially degrade tyrosine and phenylalanine. Notably, TAT and HPPD were concurrently downregulated in the high infection group, suggesting that modulation of this pathway may be a key determinant of infection outcome. Suppression of tyrosine degradation in high infection mosquitoes may inadvertently preserve critical precursors for parasite metabolism, thus facilitating parasite development.

Among the top DEGs in the low infection group was cathepsin B (CATHB), a lysosomal cysteine protease involved in protein degradation, apoptosis, and immune modulation. Its strong upregulation may reflect midgut epithelial turnover in response to microbial imbalance and infection or physical damage during parasite invasion. Another highly upregulated gene was cuticular protein 75 (CPR75), which is structurally associated with soft, flexible cuticle membranes [36]. While not previously linked to the peritrophic matrix, its expression in this context raises the possibility of a role in midgut structural remodelling. Finally, we observed a surprising downregulation of two cecropins in the low infection group. Given prior studies linking antimicrobial peptide expression and peritrophic matrix dynamics to microbiota composition [31], this pattern may signal a homeostatic shift in microbial control rather than direct immunological suppression.

### Late-stage alignment of responses and residual expression differences

By 7d pbf, gene expression patterns between high and low infection groups showed strong convergence (Pearson r = 0.45, R^2^ = 0.20, p < 0.0001; **Fig. 3D**), suggesting a return to shared physiological processes as the mosquito recovers from early infection-driven perturbations. This alignment likely reflects the completion of midgut remodelling and digestion, as well as the reduced immune and metabolic stress following parasite traversal. However, some transcriptional differences persist and may be linked to the continued presence and growth of oocysts particularly in the high infection group, where parasite burden and associated nutrient demands are greater.

Interestingly, several genes that had been downregulated in the early response phases of low infection mosquitoes, including transferrin and the two cecropins, were now upregulated. This late-stage induction of transferrin may reflect either recovery of iron homeostasis or continued attempts to restrict parasite access to this essential micronutrient [37]. Similarly, the delayed upregulation of cecropins may indicate a secondary wave of antimicrobial activity, possibly in response to shifts in microbial community structure or earlier epithelial damage.

### Microbiota composition and structure correlate with infection phenotype

To understand how the midgut microbiota may contribute to the observed *P. vivax* infection phenotypes, we examined bacterial load, diversity, and taxonomic composition in *A. darlingi* midguts across the two phenotypic groups and timepoints. Particular attention was paid to the low infection group, where early transcriptional activation of redox regulation, aromatic amino acid degradation, purine depletion, and epithelial remodelling were observed.

Total bacterial load, as quantified by 16S rRNA qRT-PCR, remained relatively stable across timepoints in low infection mosquitoes (**Fig. 4A, lower panel**). This contrasted with high infection groups, where a significant increase was detected at 1d pbf. The absence of bacterial expansion in the low infection phenotype suggests that early transcriptional responses, particularly those involving oxidative stress and metabolic restriction, may restrict microbial growth and maintain epithelial stability during a critical phase of parasite development.

**Figure 4.**
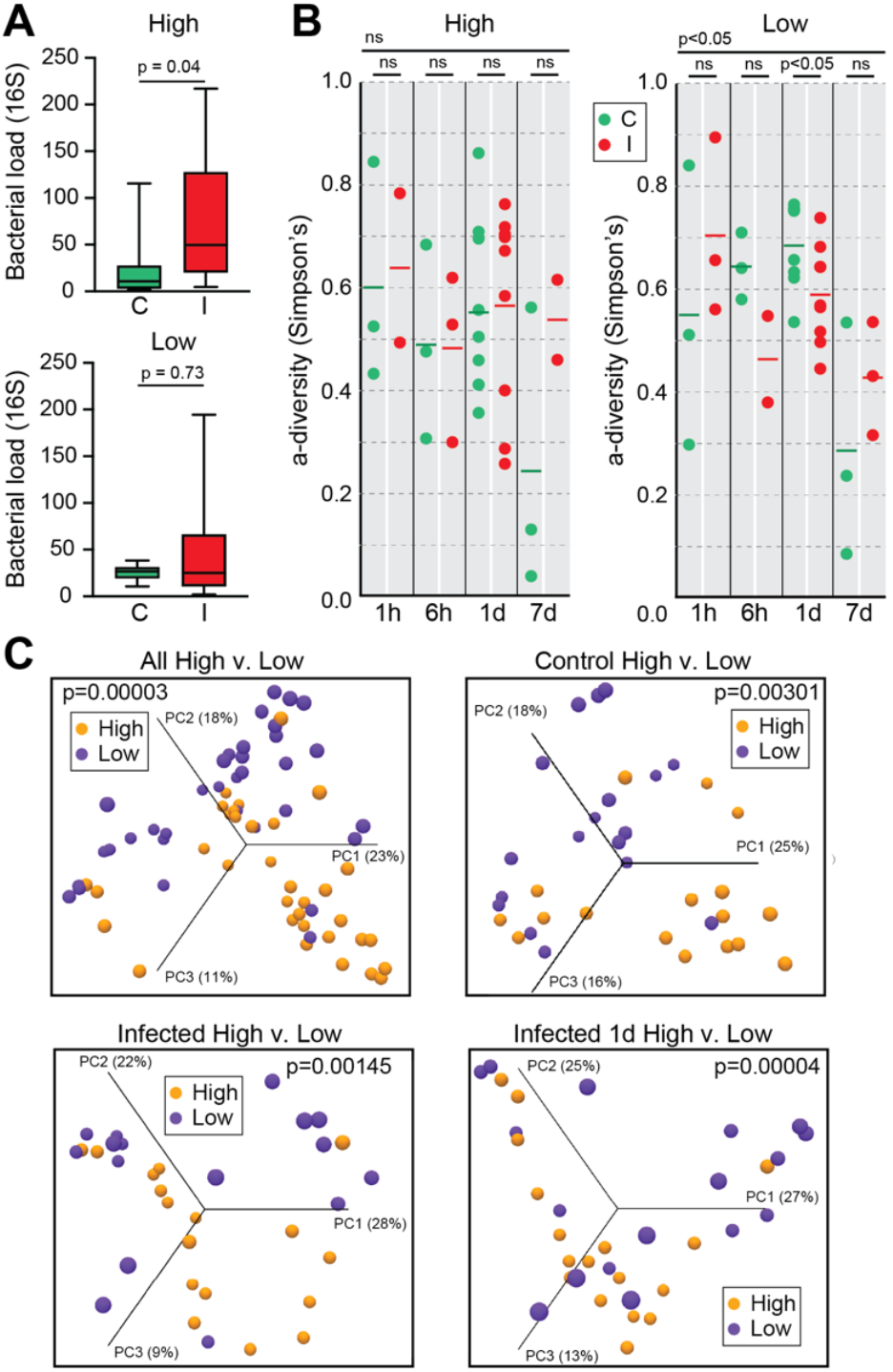
Midgut bacterial load and diversity in *A. darlingi* following *P. vivax* infection. **(A)** Total bacterial load, measured by qRT-PCR targeting the 16S rRNA gene, at 1 day post blood feeding (pbf) in high and low infection phenotypes. A significant increase in bacterial load was observed in highly infected mosquitoes (High: I) compared to controls (High: C; *p* = 0.04). No significant difference was observed in the low infection group (*p* = 0.73). **(B)** Alpha diversity (Simpson’s index) across the four timepoints pbf (1h, 6h, 1d, 7d) in high (left) and low (right) infection groups. A significant reduction in bacterial diversity was detected at 1d and 7d pbf in the low infection group (*p* < 0.05), while no significant differences were observed in the high infection group. **(C)** Beta diversity analyses using Bray-Curtis Principal Component Analysis (PCA). Top left: clustering of all samples by infection phenotype (high vs. low), showing distinct microbial community structures (*p* = 0.00003). Top right: separation of control mosquitoes only (C high vs. C low; *p* = 0.00301). Bottom left: clustering of infected mosquitoes only (I high vs. I low; *p* = 0.00145). Bottom right: strongest divergence observed at 1d pbf between high and low infected mosquitoes (*p* = 0.00004). Percent variation explained by each principal component axis is indicated.

Supporting this, alpha-diversity analysis revealed a significant reduction in microbiota diversity in the low infection group at 1d pbf (**Fig. 4B, right panel**), coinciding with peak expression of detoxification, redox, and nutrient-deprivation genes. The loss of bacterial diversity may reflect microbiota selection driven by redox or metabolic stress rather than canonical immune activity, consistent with the observed downregulation of antimicrobial peptides like cecropins.

Beta-diversity (Bray-Curtis PCA) analysis revealed consistent microbiota differences between high and low infection groups across all comparisons, including controls, suggesting that infection phenotype was associated with persistent microbiota structuring, independent of parasite exposure (**Fig. 4C**). Notably, groups producing low infection (P1, P3, P4) clustered separately from those producing high infections (P2, P5, P6), indicating that differences in host bloodmeal composition or pre-existing microbiota may act as upstream determinants of mosquito-microbiota-parasite interactions and, ultimately, infection outcome.

### Suppressive bacterial taxa and redox-linked dysbiosis in low infection mosquitoes

Taxonomic profiling at the class and order levels revealed striking differences in bacterial community composition between phenotypes (**Fig. 5A-C**). At 1d pbf, low infection mosquitoes were significantly enriched for Enterobacteriales and Pseudomonadales, two Gammaproteobacterial orders previously associated with antiplasmodial activity [38, 39]. In particular, *Pantoea, Serratia* and *Thorsellia* (Enterobacteriales), as well as *Pseudomonas* were more abundant in low infection mosquitoes (**Fig. 5B**). These genera are frequently found in wild *Anopheles* populations and include species capable of generating reactive oxygen species or producing metabolites that interfere with *Plasmodium* development, potentially linking them to the observed redox activation in low infection mosquitoes [16, 34].

**Figure 5.**
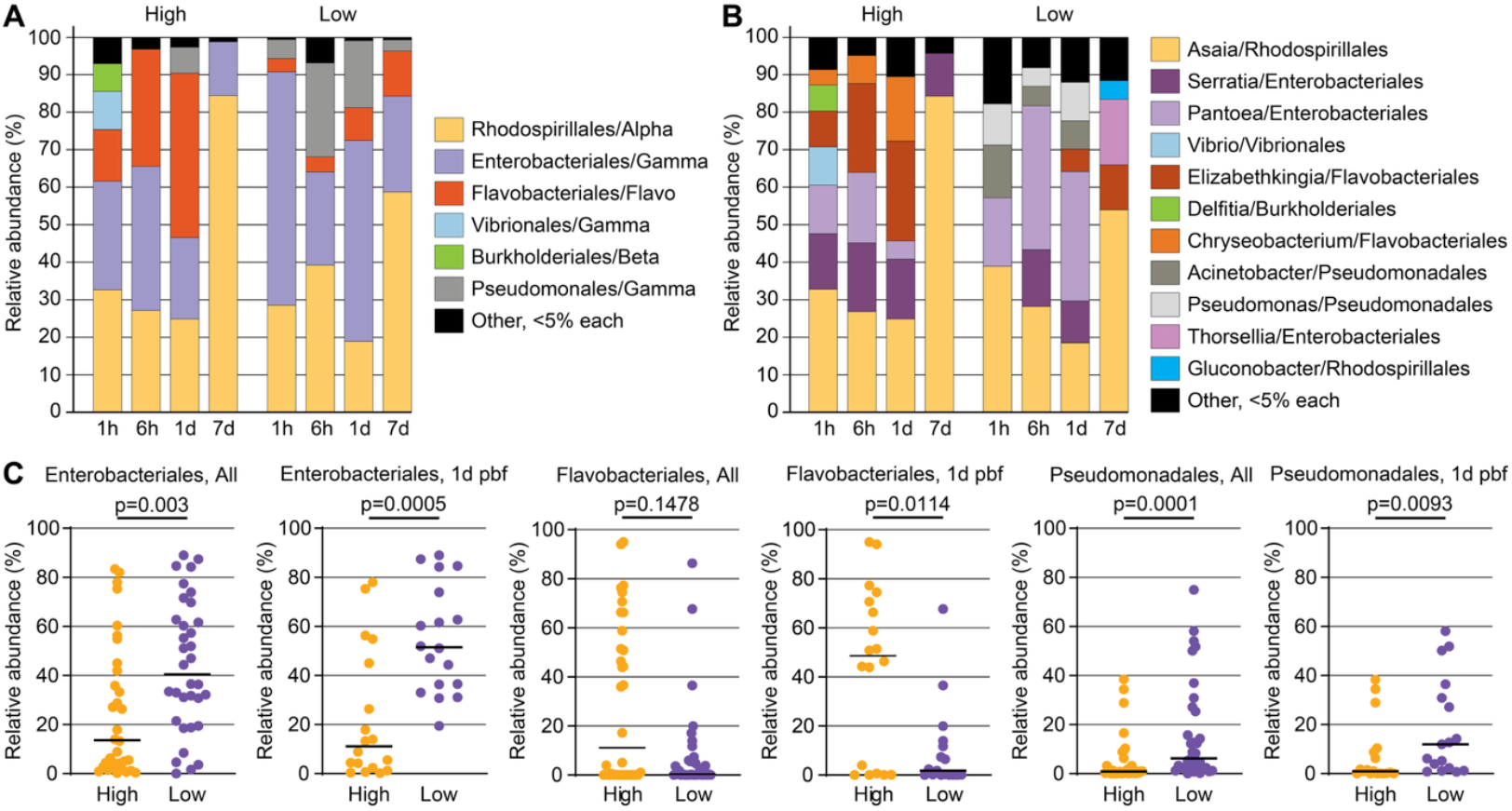
Taxonomic composition and differential abundance of midgut bacterial communities in *A. darlingi* across *P. vivax* infection phenotypes. **(A)** Relative abundance of bacterial orders across timepoints (1h, 6h, 1d, 7d post blood feeding) in high and low infection groups. Dominant orders include Enterobacteriales, Flavobacteriales, and Pseudomonadales (all Gammaproteobacteria), along with Rhodospirillales (Alphaproteobacteria) and others. **(B)** Relative abundance of representative bacterial genera within dominant orders for high and low infection groups at the same timepoints. Key differences include higher representation of *Elizabethkingia* and *Chryseobacterium* (Flavobacteriales) in high infection groups, and increased abundance of *Pantoea, Serratia, Thorsellia* (Enterobacteriales), and *Pseudomonas* and *Acinetobacter* (Pseudomonadales) in low infection groups. **(C)** Quantification of the relative abundance of key bacterial orders at all timepoints (left) and specifically at 1d pbf (right) in high vs. low infection groups. Enterobacteriales and Pseudomonadales were significantly more abundant in low infection groups (*p* = 0.003 and *p* = 0.0001, respectively), while Flavobacteriales were significantly enriched in high infection groups at 1d pbf (*p* = 0.0114). Points represent individual mosquito samples; horizontal bars indicate group medians.

In contrast, Flavobacteriales, particularly *Elizabethkingia* and *Chryseobacterium*, dominated high infection mosquitoes but were depleted in the low infection group. These bacteria have been previously associated with *A. gambiae* immune suppression and *Plasmodium* infection (Akhouayri et al., 2013; Gimonneau et al., 2014), further supporting the idea that microbiota composition may shape midgut barrier integrity and redox tone. In particular, *Elizabethkingia anophelis* has been reported to persistently colonize mosquitoes and produce factors that modulate redox balance and vector immunity and protect *Plasmodium* from human complement-mediated killing [40–42].

Taken together, these findings suggest that selective enrichment of certain bacterial taxa and reduced bacterial diversity in the low infection group reflect a microbially moderated metabolic state that is less permissive to parasite development. Rather than direct immune clearance, the low infection phenotype appears to arise from early microbiota-mediated modulation of midgut homeostasis, redox dynamics, and nutrient availability, conditions that challenge parasite survival during midgut traversal.

## Conclusion

This study provides a systems-level insight into the molecular and microbial factors shaping *P. vivax* infection outcomes in the major neotropical vector *A. darlingi*. By integrating time-resolved transcriptomic and microbiota data from experimentally infected mosquitoes, we identified two sharply divergent phenotypes, high and low oocyst burdens, which emerged despite similar infection prevalence. Importantly, these outcomes were not primarily governed by canonical immune pathways but instead arose from early and sustained interactions between midgut metabolism, microbial ecology, and bloodmeal composition.

Mosquitoes that developed low infection burdens mounted a rapid and sustained transcriptional response beginning as early as 1 hour post blood feeding, characterised by detoxification, redox regulation, aromatic amino acid catabolism, and purine depletion, likely coordinated through neurophysiological cues. These changes collectively create a nutritionally restrictive and hostile midgut environment for parasite development. Concurrently, microbial diversity contracted and became enriched in taxa such as *Pantoea, Serratia*, and *Pseudomonas*, genera previously associated with ROS production and anti-*Plasmodium* activity [43, 44]. In contrast, high-infection mosquitoes exhibited subdued metabolic reprogramming and expansion of Flavobacteriales such as *Elizabethkingia* and *Chryseobacterium*, genera linked to immune suppression and compromised epithelial integrity. These microbial shifts, along with the distinct metabolic tone of the midgut, suggest that early microbiota-mediated modulation of physiological state is a key determinant of vector competence. Our findings are consistent with and extend previous work in *Anopheles stephensi*, where *P. vivax* infection outcomes were shown to depend on specific bacterial taxa and microbiota-driven immune activation [45]. However, unlike *A. stephensi*, in which parasite clearance was primarily linked to canonical immune pathways, our results in *A. darlingi* suggest a distinct mode of vector competence regulation dominated by early metabolic and redox programming. This contrast underscores species-specific strategies of *P. vivax*–vector co-adaptation and highlights the importance of studying ecologically relevant vectors in different malaria-endemic settings.

All low infection phenotypes originated from patients with the lowest gametocyte densities. While this may at first suggest a quantitative threshold for infectivity, our findings point to a more complex model in which qualitative differences in blood composition, linked to host and parasite-derived factors, prime the mosquito midgut either for resistance or susceptibility. Such priming may be mediated by labile host metabolites or parasite signals that influence microbiota structure and midgut redox-metabolic programming, enabling the mosquito to sense and respond to the transmission potential of the bloodmeal.

Together, our findings redefine *A. darlingi* vector competence as a dynamic, emergent property of early physiological and microbial interactions rather than a fixed immune trait. This perspective opens new avenues for transmission-blocking interventions based on modulating midgut metabolism or promoting protective microbial communities. More broadly, it highlights the need to study vector– parasite–microbiota interactions in ecologically and regionally relevant systems, where distinct evolutionary and epidemiological forces shape the biology of malaria transmission.

## Methods

### Mosquito rearing

A. *darlingi* mosquitoes were obtained from a laboratory colony established at Universidad Peruana Cayetano Heredia in Iquitos, Peru [22]. The colony was maintained under standard insectary conditions at 27°C, 80% relative humidity, and a 12:12 h light/dark photoperiod. Adult mosquitoes were housed in mesh cages and provided with a 10% sucrose solution ad libitum. Female mosquitoes aged 3-5 days were used for colony propagation or experimental infections with *P. vivax*. Larvae were reared in mineral water at 28-30°C and fed Nutrafin fish food: L1/L2 instars were fed twice daily, while L3/L4 instars received three feedings per day.

### Parasitological survey

Peripheral blood samples were collected from six symptomatic malaria patients (P1-P6), aged 19 to 47 years, who presented at a local health facility in Iquitos, Peru. Thick blood smears were prepared, stained with 10% Giemsa, and examined by light microscopy for parasite detection and quantification. All patients were found to be infected exclusively with *P. vivax*. Parasite densities, including both sexual and asexual stages, were estimated using a standard assumption of 6,000 white blood cells (WBC) per μL of blood, following national guidelines established by the Peruvian Ministry of Health. Parasite counts were based on 200 WBCs, or 500 WBCs when fewer than 10 parasites were observed. Blood collection was performed with informed consent from all participants.

### Gametocytaemic blood processing and mosquito infections

Blood samples from six *P. vivax*-infected patients (P1-P6), confirmed to carry gametocytes, were processed and used to infect *A. darlingi* mosquitoes via DMFAs as described previously for *A. gambiae* [13, 26]. Briefly, patient serum was replaced with an equal volume of non-immune human serum to remove potential transmission-blocking factors. Each processed sample was then split into two aliquots: one was used directly for mosquito infection (viable gametocytes), and the other was heat-inactivated by incubation at 42°C for 12 minutes to serve as a control. Adult female mosquitoes, starved for 12 hours prior to feeding, were allowed to feed for 20 minutes on either the viable or heat-inactivated bloodmeal using glass membrane feeders. This resulted in matched infected (I) and control (C) mosquito groups for each of the six blood donors. Each infection event using a different donor blood was treated as an independent biological replicate (six replicates in total).

### Mosquito dissections and RNA isolation

*A. darlingi* mosquitoes from both *P. vivax*-infected and control (heat-inactivated) groups were cold-anaesthetised, and their midguts were dissected in phosphate-buffered saline (PBS) under a stereomicroscope at six defined timepoints pbf: 1h, 6h, 18h, 22h, 26h, and 7d. For assessment of infection intensity, an additional 10-15 midguts were dissected at 7-8 days pbf, stained with 2% mercurochrome, and examined under a light microscope for oocyst enumeration. For transcriptomic analysis, dissected midguts were immediately transferred into microcentrifuge tubes containing Trizol® reagent (Invitrogen) in pools of 30-45 midguts per timepoint and stored at −80°C. In total, 72 midgut pools were collected, representing six biological replicates (P1-P6) across two experimental conditions (infected and control) and six timepoints. Total RNA was extracted following the manufacturer’s instructions for Trizol reagent and quantified using a Qubit fluorometer (Invitrogen).

### RNAseq library preparation and sequencing

Strand-specific RNA-Seq libraries were prepared individually for each of the 72 midgut pools described above. For each sample, 0.8-2 μg of total RNA was used to generate libraries using the SureSelect Strand-Specific RNA Library Prep Kit for Illumina Multiplexed Sequencing (Agilent). Library quality was assessed using the QIAxcel Advanced System (QIAGEN) with the QIAxcel DNA High Resolution Kit (1200), and library concentrations were quantified via real-time PCR using the KAPA Library Quantification Kit (Roche). Final libraries were normalised to 2 nM and pooled for multiplexed paired-end sequencing on an Illumina NextSeq 500 platform. Raw RNA-seq data have been deposited in the European Nucleotide Archive (ENA) under the study title: RNA-Seq of *A. darlingi* infected or uninfected with *P. vivax* at various time points with high and low infections; Study ID: PRJEB29445 (ERP111747); Release Date: 03-Jan-2023.

### RNAseq data processing and analysis

The quality of paired-end sequencing reads was assessed using FastQC, and adapters and low-quality bases were trimmed using Trimmomatic v0.32.3 [46]. Cleaned reads were aligned to the *A. darlingi* genome assembly AdarC3 [47] using TopHat2 v2.0.14 with default settings [48]. Read quantification and transcript abundance were calculated as fragments per kilobase of exon per million mapped reads (FPKM) using Cufflinks v2.2.1 [49]. Differential gene expression analysis was conducted with Cuffdiff v2.2.1 [50], comparing *P. vivax*-infected mosquito midguts to their corresponding heat-inactivated controls at each dissection timepoint. To assess the consistency among replicates and overall transcriptional patterns, unsupervised hierarchical clustering was performed using the ward.D2 method on global transcript profiles. Clustering and heatmap visualization were performed with Cluster 3.0 and Java TreeView [51], respectively. Pearson correlation analyses of log2 fold-change profiles between high and low infection groups were performed to evaluate the temporal similarity in transcriptional responses across timepoints.

### Gene Ontology term assignment and enrichment

GO terms were assigned to *A. darlingi* transcripts (AdarC3.5 gene set, release date: June 2017) by orthology-based mapping to the *A. gambiae* gene set (AgamP4.6) using BioMart. Functional enrichment analysis of differentially expressed genes (DEGs) was conducted using the PANTHER classification system available through VectorBase. Enrichment was assessed across the three GO categories: Biological Process (BP), Molecular Function (MF), and Cellular Component (CC), with significance determined by Fisher’s exact test and false discovery rate (FDR) correction (Padj < 0.05).

### Quantification of *A. darlingi* gut microbiota by real-time qPCR

Quantitative real-time PCR (RT-qPCR) was used to assess bacterial load in the midguts of *P. vivax*-infected and control *A. darlingi* mosquitoes by targeting the bacterial 16S ribosomal RNA gene (16S rRNA) as described [52]. Total RNA (1 µg) from each of the 72 midgut pools was treated with DNase I (Invitrogen™) at 37 °C for 2 hours and reverse-transcribed using the RevertAid First Strand cDNA Synthesis Kit (Thermo Scientific™), following the manufacturer’s protocol. RT-qPCR reactions were performed in triplicate using SYBR Green PCR Master Mix (Applied Biosystems™) in a final volume of 20 µL. Expression of the 16S rRNA gene was normalised against the *A. darlingi* reference gene Rp49 (ADAC007403) using the 2^–^ΔCT^ method [53]. RT-qPCR primers for *A. darlingi* Rp49 gene expression analysis were designed using Primer 3 online tool and their efficiency was determined by RT-qPCR: Rp49-F, ACAGTACCTGATGCCGAACA; Rp49-R: TTCTGCATCATCAGCACCTC Statistical significance of microbiota load differences between infected and control groups at the 1d pbf timepoint of midgut invasion was assessed using the Mann-Whitney U test.

### Multiplex bacterial *16S rRNA* amplicon Illumina sequencing

The gut bacterial communities of *P. vivax*-infected and control *A. darlingi* mosquitoes were profiled using multiplex 16S rRNA amplicon sequencing as described [54]. Amplicon libraries targeting the V4 hypervariable region of the bacterial 16S rRNA gene were generated via PCR using cDNA synthesised from previously DNase-treated midgut RNA as template. Each reaction used a single universal forward primer and a set of reverse primers tagged with unique barcodes (indexes) to enable sample multiplexing. PCRs were performed in triplicate per sample, and resulting amplicons were pooled and purified using AMPure XP magnetic beads (Beckman Coulter). Purified libraries were quantified with the KAPA Library Quantification Kit (Roche®), normalised to 2 nM, and pooled equimolarly. Sequencing was performed using a V2 MiSeq Reagent Kit on the Illumina MiSeq platform.

### *16S rRNA* amplicon Illumina sequencing analysis

Raw sequencing data were first processed using the MiSeq Reporter software (Illumina) to remove adapter sequences and generate demultiplexed FASTQ files. Subsequent analysis was carried out using the Microbial Genomics Module of CLC Genomics Workbench v10.1.2 (QIAGEN), following the manufacturer’s standard pipeline. Paired-end reads were merged using the following parameters: minimum score of 8, gap cost of 3, mismatch cost of 2, and no maximum unaligned bases. Merged reads were trimmed to 264 bp, and samples with fewer than 100 reads or below 50% of the median read count were excluded. Operational Taxonomic Unit (OTU) classification was performed using the SILVA v119 16S rRNA reference database with a minimum sequence identity threshold of 97%. Low-abundance OTUs representing <0.05% of total reads within a sample were filtered out. Microbial diversity was assessed using both alpha and beta diversity metrics. Alpha diversity was calculated using Simpson’s diversity index, which incorporates both OTU richness and evenness, placing greater weight on dominant taxa. Beta diversity was assessed using Bray-Curtis dissimilarity distances and visualised via principal coordinates analysis (PCoA) using the top three principal components. Clustering patterns were statistically evaluated using PERMANOVA, based on sample metadata.

## Ethics statement

All procedures involving *A. darlingi* infections with naturally circulating *P. vivax* gametocytes from human carriers were conducted at Universidad Peruana Cayetano Heredia under approved ethical guidelines. Ethical approval was granted by the institutional review board of Universidad Peruana Cayetano Heredia (Protocol No. R157-13-14). Informed consent was obtained from all participating individuals prior to blood collection, in accordance with national and institutional ethical standards.

## Acknowledgements

We thank Lutecio Torres, Gerson Guedez, Cristian Rodriguez, Juan Michi and Zaira Villa for their assistance with the DMFAs, and Daniel Lawson for help with genome alignments and de novo gene model generation. We are grateful to the six volunteer patients who donated blood for this study. This work was supported by a Young Investigator Award to J.A.S.N. from the São Paulo Research Foundation (FAPESP; grant number 2013/11343-6), a mobility grant to J.A.S.N. and G.K.C. under the FAPESP-Imperial College London cooperative agreement (grant number 2014/50454-0), and a Special Visiting Fellowship from the National Council for Scientific and Technological Development (CNPq; grant number 401433/2014-5) to J.A.S.N. and G.K.C. under the Science without Borders Program. B.C.C. was supported by a CNPq postdoctoral fellowship (grant number 154727/2016-4). G.C.K. and D.V. were also supported by a Wellcome Trust Investigator Award (grant number 107983/Z/15/Z) and a Medical Research Council (MRC) project grant (MR/T000929/1). C.T.R. and M.M. were supported by the National Institutes of Health-National Institute of Allergy and Infectious Diseases (NIH-NIAID) ICEMR-Amazonia program U19AI089681. We are grateful to Jan E. Conn and Joseph M. Vinetz for facilitating the collaboration with ICEMR-Amazonia and enabling this study.

## Author Contributions

Conceptualization: D.V., G.K.C. and J.A.S.N.; Methodology: M.M., D.V., G.K.C. and J.A.S.N.; Formal analysis: B.C.C., P.H.A.A., A.J., D.P.A., B.M, D.V., G.K.C. and J.A.S.N.; Investigation: B.C.C., K.V., P.H.A.A., B.T.N., D.P.A., D.V., G.K.C. and J.A.S.N.; Resources: M.M., D.V., G.K.C. and J.A.S.N.; Data Curation: A.J. and B.M.; Writing - Original Draft: B.C.C., D.V., G.K.C. and J.A.S.N.; Writing - Review & Editing: D.V. and G.K.C.; Visualization: G.K.C.; Supervision: D.V., G.K.C. and J.A.S.N.; Project administration: M.M., D.V., G.K.C. and J.A.S.N.; Funding acquisition: D.V., G.K.C. and J.A.S.N.

## Materials & Correspondence

All correspondence and materials requests should be addressed to G.K.C. or J.A.S.N. or D.V.

## Supplementary Datasets

**Dataset 1. Parasitological study and oocyst load and prevalence data**.

**Dataset 2. Differential gene expression data in high infection groups**.

**Dataset 3. Differential gene expression data in low infection groups**.

**Dataset 4. Gene Ontology (GO) enrichment data in high and low infection groups**.

**Dataset 5. Differentially expressed genes at each timepoint**.

